# Progression Analysis of Disease with Survival (PAD-S) by SurvMap identifies different prognostic subgroups of breast cancer in a large combined set of transcriptomics and methylation studies

**DOI:** 10.1101/2022.09.08.507080

**Authors:** Jaume Forés-Martos, Beatriz Suay-García, Raquel Bosch-Romeu, Maria Carmen Sanfeliu-Alonso, Antonio Falcó, Joan Climent

## Abstract

Progression analysis of disease (PAD) is a methodology that incorporates the output of Disease-Specific Genomic Analyses (DSGA) to an unsupervised classification scheme based on Topological Data Analysis (TDA). PAD makes use of data derived from healthy individuals to split individual diseased samples into healthy and disease components. Then, the shape characteristics of the disease component are extracted trough the generation of a combinatioral graph by means of the Mapper algorithm. In this paper we introduce a new filtering function for the Mapper algorithm that naturally integrates information on genes linked to disease-free or overall survival. We propose a new PAD-extended methodology termed Progression Analysis of Disease with Survival (PAD-S) and implement it in an R package called SurvMap which allows users to carry out all the steps involved in PAD-S, as well as in traditional PAD analyses. We tested PAD-S methodology using SurvMap on a large combined transcriptomics breast cancer dataset demonstrating its capacity to identify sets of samples displaying highly significant differences in terms of disease free survival (*p* = 8 *×* 10^−14^) and idiosyncratic biological features. PAD-S and SurvMap were also able to identify sets of samples with significantly different relapse-free survivals and molecular profiles inside breast cancer intrinsic subgroups (luminal A, luminal B, Her2, and basal). Finally, to illustrate that PAD-S and SurvMap are general-purpose analysis tools that can be applied to different types of omics data, we also carried out analyses in a breast cancer methylation dataset derived from The Cancer Genome Atlas (TCGA) identifying groups of patients with significant differences in terms of overall survival and methylation profiles.

## 1 Introduction

Topological data analysis (TDA) is an emerging field in the context of biological and biomedical research. The two major analysis tools derived from TDA are persistent homology and mapper. Persistent homology borrows ideas from abstract algebra to identify particular aspects related to the shape of the data such as the number of connected components and the presence of higher-dimensional holes, whereas mapper condenses the information of high-dimensional datasets into a combinatory graph or simplicial complex that is referred to as the skeleton of the dataset. Both approaches have been successfully applied to the study of biological and biomedical datasets including disease diagnosis, subtyping, and staging, omics data analysis including cell-type classification in single-cell RNAseq studies, and macromolecule structure analysis, among others [1].

One early contribution to the use of mapper in the context of omics data analysis was provided by Nicolau and co-workers [2]. In their work they implemented a method termed progression analysis of disease (PAD) which combined the use of transcriptomics data transformed using Disease-Specific Genomic Analysis (DSGA) with mapper to identify subsets of breast cancer patients [2].

DSGA was first proposed by Nicolau and co-workers as a method to apply before clustering and class prediction analyses that resulted in an increase in their performance [3]. DSGA measures the amount of deviation that disease samples present when compared to matched healthy control tissues, isolating and separating the disease-component (*D*_*c*_) from the normal-component (*N*_*c*_) portion of the data. DSGA first creates a model for normal data defined as a linear subspace of the normal tissue samples which is termed as Healthy State Model (HSM). Then, the *D*_*c*_ of each diseased sample is extracted by computing its deviation from the HSM in terms of the residuals of a linear model fit. This method was found to perform better than traditional log-ratio transformation in class prediction by placing tumor samples in classes defined by clinical and pathological co-variates with better error rates [3]. In addition, DSGA was also found to be able to uncover new biological aspects of the input data in breast cancer samples [3]. Although DSGA was first used in the context of transcriptomics datasets, it can be easily generalized to other high-dimensional omics data types such as methylation or proteomics.

DSGA was later extended to *Progression Analysis of Disease* (PAD) [2]. In PAD high dimensional data previously transformed using DSGA is used as an input for the Mapper algorithm [4], which builds a geometric representation of the data using preassigned guiding functions called filters. In PAD analyses, the output is a simple low-dimensional shape that can be visualized as a graph in which nodes typically represent groups of samples and edges connect nodes that share samples. PAD has been successfully applied to the analysis of a breast cancer transcriptomic dataset allowing the identification of a subgroup of *Estrogen Receptor-positive* (*ER*^+^) breast cancer samples displaying diverse molecular characteristics and 100% survival rates [2].

Here we present Progression Analysis of Disease with Survival (PAD-S), an extension of PAD that allows for a natural integration of prior information on genes linked to survival in the Mapper filtering function used for PAD analyses. This guiding function unravels data according to the extent to which each sample deviates from the HSM in a way in which the deviation of each gene is weighted by its degree of association with overall or relapse-free survival. In addition, PAD-S also implements a new function for the HSM generation that, in contrast with the previously employed function, which was based on the Wold’s goodness of fit measure [5], does not require any user’s input. The new approach is based on an automatic optimal singular value hard threshold identification method for rectangular matrices described in [6]. In addition, a new criteria based on Silhouette analysis [7] for the identification of the optimal number of clusters has also been introduced. All functions and methods necessary to carry out both PAD-S, as well as traditional PAD analyses have been implemented and wrapped into SurvMap, an R package available at (https://github.com/jfores/SurvMap). We tested PAD-S and SurvMap on a large combined array-based breast cancer (BC) transcriptomics dataset that included more than 4200 samples. PAD-S and SurvMap were able to identify sets of patients displaying significant differences in terms of disease-free survival and idiosyncratic molecular profiles. In addition, we were also able to identify subgroups of better and worse prognosis inside BC intrinsic subtypes (luminal A, luminal B, Her2, and basal). Finally, To highlight the PAD-S and SurvMap capacity to work using other omic data types, we carried out analyses employing breast cancer methylation data derived from TCGA and identified sets of patients with diverse methylation profiles and significantly different prognosis in terms of overall survival.

## 2 Material and Methods

### 2.1 Breast cancer combined dataset construction

We carried out searches in Gene Expression Omnibus (GEO) [8] for array-based gene expression datasets including breast cancer and healthy breast tissue samples. To apply homogeneous reprocessing methods and to ensure integration, only two Affymetrix array platforms (hgu133A and hgu133plus2) were selected for the analysis. Datasets including BC samples were required to present clinical information about relapse-free survival. When available, we extracted information about the following co-variates: Age at diagnosis, tumor size, tumor grade, estrogen receptor (ER) status, progesterone receptor (PR) status, and human epidermal growth factor receptor 2 (HER2), and the presence of affected lymphatic nodes, as well as, relapse-free-survival and overall survival information.

Each dataset was independently pre-processed as follows: First, raw cell files were loaded into R’s environment using the affy package [9]. The arrayQualityMetrics package was used to detect potential outlier samples [10]. Three different quality metrics were computed, the first based on between-array distances, the second on the comparison of the array intensity distributions, and the third on the inspection of MA plots. Those samples targeted as potential outliers by the three methods were excluded from the analysis. When both tumour and control samples were present in the same study, the quality control step was carried out independently for tumoral samples and controls. This prevents the preferential targeting of samples belonging to a specific group when the study design is unbalanced. After quality control and outlier samples removal, background correction, summarization, and quantile normalization were carried out using the fRMA method implemented in the R package fRMA package [11].

To avoid the inclusion of duplicated samples in the final dataset, we made use of the DupChecker package [12]. For each CEL file, DupChecker generates an MD5 fingerprint that can be used to identify redundant samples. Samples presenting redundant MD5 fingerprints were removed by randomly selecting one of them prior to dataset combination.

All individual studies were then merged in a single matrix, based on the set of 22277 shared probes present in both microarray platforms. Individual datasets were concatenated by columns. Batch effect correction was then s carried out using the Combat function from the sva package [13]. Finally, we filtered probes that did not target genes with valid gene id and collapse those probes targeting the same gene by thanking those presenting the highest row variance using the collapseRows function from the WGCNA package [14, 15]. The final dataset contained 4360 samples and 12399 genes. The full set of scripts used to construct the dataset, as well as all the analyses carried out in this work can be found in Supplementary File 1.

### 2.2 Methylation data obtention and preprocessing

Breast cancer methylation data was downloaded and pre-processed using the TCGABiolinks package [16]. The Beta values for a total of 485577 probes and 895 samples were obtained using the GDCquery, GDCdownload, and GDCprepare functions. Probes presenting missing values in more than 50% of the samples were filtered out, leaving a total of 395777 probes. The remaining missing values were imputed using the impute.knn function from the impute package. [17].

### 2.3 Disease-Specific Genomic Analysis (DSGA)

The first step in PAD-S analysis is to transform the original data using the DSGA method. DSGA measures the amount of deviation that a diseased sample presents when compared to matched healthy control tissues, isolating and separating the disease-component (*D*_*c*_) from the normal-component (*N*_*c*_) portion of the data. It involves three steps: (*i*) Flat construction. (*ii*) HSM construction, and (*iii*) Disease component obtention for the full dataset.

#### 2.3.1 Flat construction

In this step, we only make use of the healthy or normal tissue samples available in our dataset. The idea behind flat construction is to reduce or to smooth aspects of the data that are idiosyncratic to each normal or healthy tissue sample. Let’s denote *N* as the matrix containing the normal tissue samples of our dataset organized by columns. In the case of transcriptomics data, we start from the *R* normal tissue gene expression vectors. *N*_1_, *N*_2_, …, *N*_*R*_. Next, we compute the flattened vectors 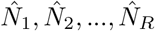. The flattened vector 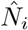 is obtained by fitting a 0-intercept least-squares linear model from all *N*_*i*_-normal tissue vectors *N*_1_, *N*_2_, …, *N*_*i*−1_, *N*_*i*+1_, …, *N*_*R*_ as predictor variables, that is,

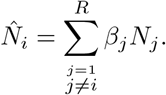

Here 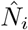 is the *i*-th flat vector and *β*_*j*_ is the coefficient associated with normal tissue sample *N*_*j*_. Then after carrying out this step separately for each healthy tissue sample we obtain the flat matrix in which each column represents a flatted healthy tissue sample:

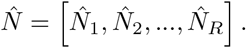

#### 2.3.2 Healthy State Model (*HSM*) generation

The flat matrix 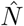 is then used to compute the HSM. In traditional PAD singular value decomposition followed by Wold’s invariant [5] computation was employed to select the most informative number of singular values. This method involved the visual inspection of a graph plotting the Wold’s invariant against the dimension of the data by the user and the manual selection of the appropriate dimension based on the presence of spikes or jumps on it. (See references [2] and [3]). In contrast, we implemented an automatic approach based on a reported method for rectangular matrix de-noising using singular value thresholds [6].

The idea behind the method is that our flat data matrix 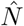 can be expressed as the sum of a signal matrix and a noise matrix.

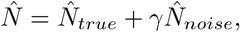

Where 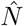 is the flat matrix, 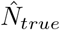 is an underlying low-rank matrix that contains the true signal and 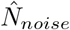 is a noise matrix in which entries are assumed to be a sample of i.i.d. random variables extracted from a Gaussian distribution, with zero mean and unit variance. Finally, *γ* is a parameter that indicates the magnitude of the noise. Let’s denote the singular value decomposition of our flat matrix as:

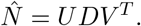

Where *D* is a diagonal matrix in which the elements of the diagonal are the singular values of 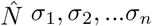, with *n* being the number of columns of 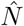, ordered from high to low and *U* and *V* ^*T*^ are matrices containing the left and right singular vectors in columns and rows, respectively.

In the case of rectangular matrices with unknown *γ* the optimal hard threshold for singular value selection is:

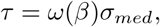

where *σ*_*med*_, is the median of the values {*σ*_1_, *σ*_2_, …*σ*_*n*_} and *ω*(*β*) is defined by

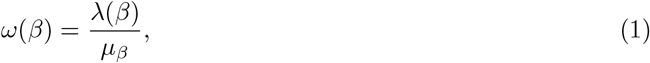

and where:

a. The numerator *λ*(*β*) in (1) is obtained through the following expression:

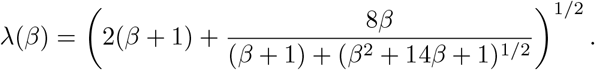

Here *β* is the aspect ratio of our input matrix 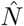, that is,

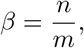

with *m* and *n* being the number of columns and rows of the matrix 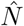, respectively.
b. The denominator *µ*_*β*_ in (1) is found by computing numerically the upper bound of the following definite integral in the range [(1 − *β*)^2^, (1 + *β*)^2^]:

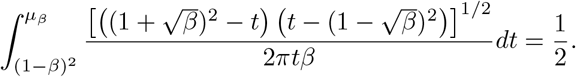

Only those singular values larger than the obtained threshold will be kept.

The new de-noised lower dimensional matrix is termed as the Healthy State Model (*HSM*). To construct it assume we need a diagonal matrix termed 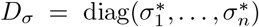 which is obtained from *D* = diag(*σ*_1_, …, *σ*_*n*_) by using that

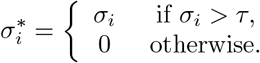

Now, the *HSM* is defined as

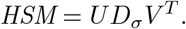

#### 2.3.3 Disease component computation

The final step in DSGA transformation consists on the computation of the disease vectors for each sample in the original dataset. Let’s denote our original matrix including both healthy and disease samples as *O*. Then, we fit a zero intercept linear model for each data vector *O*_*i*_ present in *O* to the columns of the *HSM* -matrix, that is,

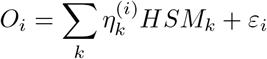

Here 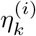 are the coefficients obtained from the best approximation of the column space of the matrix *HSM* to *O*_*i*_ and *ε*_*i*_ is its corresponding residual. Now, considering the matrix

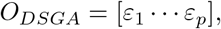

we remove the portion of each sample that best mimics the expression patterns of healthy tissues and remain with the vector of residuals that carry out the information about how much each gene of a particular sample deviates from the values observed in healthy tissues.

### 2.4 Feature Selection

Once the disease component matrix *O*_*DSGA*_ has been constructed, the next step consists of the selection of features (Genes, Proteins, Methylated CpGs, …) for downstream analysis. The feature selection procedure is based on both the variability of each probe across the disease component matrix and the degree of association of each probe with either disease-free or overall survival.

Associations between the levels of each feature (i.e. expression, methylation) with survival are computed using univariate Cox proportional hazard models using the original data matrix. The *Z*-scores derived from the Cox proportional hazard models fits representing the degree of association of each feature with survival are then stored in the vector *Z*_*cox*_. Then, the standard deviation across samples of each feature is computed using the *O*_*DSGA*_ matrix. The vector of standard deviations is then stored as *O*_*sd*_. To avoid values between 1 and −1, +1 is added to all positive values in vectors *Z*_*cox*_ and *O*_*sd*_, and −1 is added to all negative values of both vectors. The element-wise product between *Z*_*cox*_ and *O*_*sd*_ matrices is taken and the top and bottom *n* features of the distribution of the product are selected for further analysis.

Then, the *O*_*DSGA*_ matrix is filtered keeping the selected features. This matrix will constitute the input for the mapper algorithm.

SurvMap also implements the traditional PAD criteria for feature selection, in which for each DSGA-transformed feature, the absolute values of the 5-th and the 95-th percentiles are computed and the larger is kept. A user-defined percentile threshold is then applied to select the top genes of the previous distribution.

### 2.5 Mapper

Mapper is a mathematical tool based on topological data analysis that is robust to noise and changes in notions of distance and similarity. The output of Mapper is a combinatorial graph with a multi-resolution form, a characteristic that can be used to distinguish between real features and artifacts in the data [4]. It can work with diverse measures of similarity other than euclidean distances including correlations. The method starts with a filtering function *f* that provides a map from the multidimensional dataset to R. The range of the *f* function is then divided into overlapping pieces. Next, data points placed inside the range of each overlapping fragment are clustered. Each cluster can be viewed as a bin of data points. Edges connecting bins are added if a particular pair of bins share data points in common. The next sections describe each step in more detail.

#### 2.5.1 Filtering function in PAD and PAD-S

The first step in Mapper consist in the choice of an appropriate filter function. A filter function *f* maps each point in our dataset denoted by *X* to ℝ.

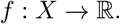

Several different functions can be chosen to act as filters and the output of Mapper will be dependent on this choice. These functions include density estimators, eccentricity, and graph Laplacian [4]. In PAD analysis the filter function of choice was the vector of magnitude in the *L*^*p*^ norm, as well as *k* powers of this magnitude.

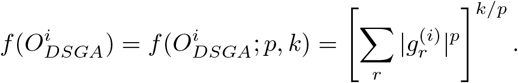

Here 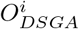 denotes the *i*-th column vector of our disease component matrix, which contains the values of each selected feature in this sample with coordinates 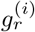. Note that if *k* = 1 and *p* = 2, the function simply computes the standard (Euclidean) vector magnitude of each column.

In the case of PAD and Mapper, the filtering function measures the overall amount of deviation from the null hypothesis (Healthy State Model) for each sample. The function is large when a large number of genes deviate from the expression levels of the HSM either in positive or negative directions. The choice of larger values of *p* implies that the weight of genes with larger expression levels will be grater. The distance function used for mapper was the correlation distance.

In the case of PAD-S and SurvMap a variant of this filtering function which takes into account the magnitude of the association between a particular feature and disease-free or overall survival is implemented. In particular the filter function is defined as follows:

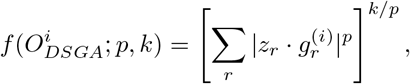

where *z*_*r*_ is the *z*-value derived from the Cox proportional hazard models analysis and 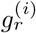 the *i*-th disease component value for *r*-th feature.

Note that in this case, to avoid values between 1 and −1, +1 is added to all positive values in vectors *Z*_*cox*_ and 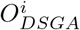, and −1 is added to all negative values of both vectors. Therefore, the amount of deviation of each feature to the *HSM* is multiplied by the degree of association of this particular feature with survival, and the initial overlapping sets of samples incorporate this information.

#### 2.5.2 The One Dimensional Mapper Algorithm

Mapper will be applied on *O*_*DSGA*_ by using one of the filter functions defined in the previous section.

In addition, Mapper requires the use of a specific distance metric. To this end, SurvMap allows to choose among different distance metrics including correlation and euclidean distances.

Once the overlapping sets have been generated by the filtering function, the samples included in each fragment are clustered by choosing one of the following methods: single linkage, average linkage, complete linkage, or k-medioids. Note that in the original Mapper article [4] no restrictions are imposed about type of clustering method to be used.

The output graph *G* = *G*(*V, E*) is then defined putting each cluster as a node (or vertex) of the nodes that share samples are connected with an edge.

#### 2.5.3 Methods for the identification of the optimal number of clusters

The original algorithm employed for the selection of the optimal number of clusters proposed in the Mapper seminal article [4] is also used in classical PAD analysis [2]. It is based on the construction of an histogram of *k* intervals from the vector of the cluster edge lengths, denoted by *C*. Empirically, shorter edges are required to connect points within a particular cluster, compared to those required to join two different clusters. Therefore looking at the histogram of edge lengths generated in a particular node, shorter edges will connect points within each cluster. Those will have a smooth distribution in the histogram. In contrast, edges that are required to merge the different clusters will be disjoint from the smooth distribution. Generating the histogram using *k* intervals will generate a set of empty intervals after which the edges which are required to merge clusters will appear. Then, by selecting edge lengths shorter that those at which the empty interval is observed we can recover a clustering of the date. Increasing the value of *k* will increase the number of recovered clusters whereas reducing it will decrease it.

An additional method based on the analysis of Silhouettes [7] has been implemented to find the optimal number of clusters in PAD-S and SurvMap. To this end, for a particular data point *x* included in cluster *C*_*i*_ we first define *a*(*x*) as the average distance of data point *x* to all other points *y* in the cluster. Thus,

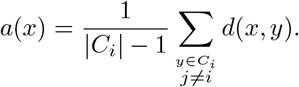

Then, *b*(*x*) is defined as the minimum mean difference of point *x* to any other cluster *C* of which *x* is not a member, as the average distance from *x* to all points *y* in *C* where *C* ≠ *C*_*i*_.

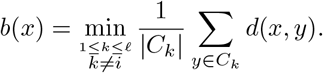

The cluster which has the minimum average distance to point *x* is said to be the neighbor cluster of *x*.

Now, the concept of Silhouette for point *x* can be defined as:

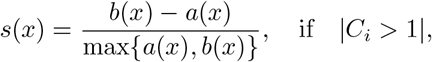

that is if the number of elements in the cluster is larger than 1. When a particular cluster *C*_*i*_ contains only a single object, it is unclear how *a*(*x*) should be defined, and therefore we set the silhouette to 0.

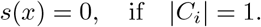

The average of the *s*(*x*) values of all points grouped in a particular cluster *C*_*i*_, i.e.

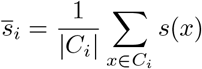

indicates how well grouped are the members of this cluster, whereas the average of all data points

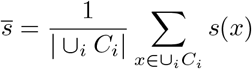

indicates how well the available data points have been clustered in general.

To select the optimum number of clusters inside each node, the average silhouette values 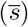 are computed for all the possible partitions from 2 to *n* – 1, where *n* is the number of samples inside a specific node. Then the *n* that produces the highest value of 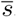. and that exceeds a specific threshold of 0.25 is selected as the optimum number of clusters. Otherwise, all samples are assigned to a unique cluster.

### 2.6 Assignment of promiscuous samples to individual nodes

The combinatorial graph produced by PAD-S and SurvMap contains nodes that are joined by an edge if they include shared data points or samples. For subsequent analysis such as survival, differential methylation or gene expression it is important to place promiscuous samples in individual nodes. We devised a method to accomplish this task. To this end we write two generic nodes *V*_*α*_ and *V*_*β*_ that are sets including vectors extracted from matrix *O*_*DSGA*_, as matrices

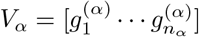

and

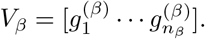

Moreover, we assume the existence of a vector

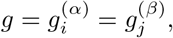

shared by both matrices as column. To decide whether assign *g* either to *V*_*α*_ or *V*_*β*_ we proceed as follows. We first construct the matrices 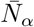, and 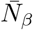 as:

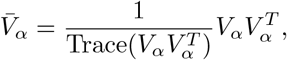

and

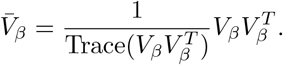

Observe that, now 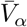 and 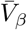 are square matrices of the same size, symmetric, semi-definite positive and with unitary trace.

Next, we compute the ratios

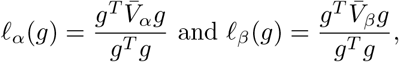

where *g*^*T*^ denotes the transpose of vector *g*. It can be shown that

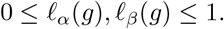

The decision criteria is: *g* ∈ *V*_*α*_ if and only *ℓ*_*α*_(*g*) *> ℓ*_*β*_(*g*). Otherwise *g* ∈ *V*_*β*_.

### 2.7 Differential gene expression and individual sample pathway enrichment analysis methods

Differential gene expression analyses across nodes were carried out employing the original gene expression matrices and performing two-tailed Wilcoxon tests comparing those samples placed in a particular node to all other samples with the help of the GSALightning package [18]. Probes presenting adjusted p-values lower than 0.05 were considered to be differentially expressed or methylated. To determine if the breast cancer nodes identified by PAD-S and SurvMap presented alterations in different pathways and to easily visualize those changes across the sequence of breast cancer nodes we used the original gene expression matrix and the Gene Set Variation Analysis (GSVA) package [19]. This package allows the user transform the gene expression matrix into a gene set enrichment score matrix employing the GSVA method. The Hallmarks of cancer gene set was employed for the analyses.

### 2.8 Weighted Gene Co-Expression Networks construction

We analysed if specific gene co-expression modules were associated with some of the breast cancer patient subgroups determined by PAD-S using Weighted Gene Co-expression Network Analysis [14, 15]. In short, starting from the original gene expression dataset, we carried out WGCNA consensus module detection by constructing consensus signed hybrid networks using bicor as a distance function.

## 3 Results

### 3.1 SurvMap package outline

SurvMap implements all necessary functions to perform Progression Analysis of Disease with Survival (PAD-S), as well as several other additional functions aimed to help with the visualization of the results and designed to carry out diverse analysis types using PAD-S output such as differential gene expression and survival analyses between the groups identified by SurvMap as well as individual sample enrichment analysis. Table 1 shows the most relevant core and auxiliary SurvMap functions.

**Table 1:** Table showing the core and auxiliary functions implemented in the SurvMap package.

|  |  |
| --- | --- |
| **Core Functions** |  |
| **flatten_normal_tiss()** | Produces the flattened normal tissues matrix from the original samples derived from healthy tissues. |
| **denoise_rectangular_matrix()** | Removes Gaussian noise from the flattened healthy tissue matrix. |
| **generate_disease_component()** | Extracts the diseases component of each sample. |
| **cox_all_genes()** | Computes the association of expression levels with relapse-free survival or overall survival for each gene. |
| **gene_selection()** | Selects features using PAD original criterion. |
| **gene_selection_surv()** | Selects features based in the new PAD-S feature selection criterion. |
| **lp_norm_k_powers()** | Filtering function to carry traditional PAD. |
| **lp_norm_k_powers_surv()** | Filtering function to carry out PAD-S. |
| **one_D_Mapper()** | Runs the one-dimensional mapper algorithm. |
| **Auxiliary Functions** |  |
| **plot_mapper()** | Plots the output combinatorial graph produced by the one_D_Mapper function. |
| **assign_promiscuous_samples()** | Assigns samples placed in more than one node to individual nodes. |
| **survival_analysis_multiple_groups()** | Performs survival analysis (log-rank tests) based on the placement of the samples in specific nodes and produces Kaplan-Meier plots. |
| **perform_wilcoxon_each()** | Carries out differential gene expression (Wilcoxon tests) comparing those samples placed into a particular to all other samples. |
| **select_top_diff_genes_groups()** | Selects the $n$ top differentially expressed genes in a particular node. |
| **plot_heatmap_data()** | Produces a heatmap based on the expression values of the top differentially expressed probes. |
| **plot_genes_by_groups()** | Creates violin plot visualizations of the levels of specific probes in each identified node. |
| **perform_GSVA_dataset()** | Carries out GSVA analyses for the complete dataset based on sample-node assignments. |
| **plot_GSVA()** | Produces a heatmap displaying the results generated by the perform_GSVA_dataset() function. |

### 3.2 Breast Cancer Combined dataset analysis results

#### 3.2.1 Testing the effect of Parameters on SurvMap’s output

We first constructed a combined breast cancer gene expression dataset following the procedures described in the material and methods. The combined dataset included a total of 4360 samples divided into 4219 breast cancer samples and 141 breast healthy tissue samples obtained from 25 different array-based transcriptomics studies. Next, we divided the combined dataset into two blocks of approximately the same size by randomly selecting the same number of cases and controls. Supplementary Table 1 shows the studies included to construct the dataset, the number of samples, and the placement of each sample in each subset (A or B).

Subset A incorporated 2173 samples divided into 2104 cases and 69 controls. We employed the SurvMap package to carry out progression analysis of disease with survival using Subset A. The healthy state model was computed using the 69 healthy breast tissue samples which was followed by the computation of the disease component. Associations between the expression levels of each gene with disease-free survival were measured by fitting Cox proportional hazard models. Supplementary File 2 shows the association between gene expression levels and survival for all tested genes in Subset A. Feature selection was carried out employing the **gene selection surv()** function and the top 200 features were selected for downstream analyses.

We studied the effect of a range of parameter values in the outputs of PAD-S and SurMap, including the number of initial intervals 5 − 25, the overlap percentages: 0 −0.9, and the number of bins 5 − 20, and analyzed their influence in the final number of nodes identified by mapper, their average content in samples of nodes, and the number of ramifications present in the combinatorial graph.

The initial number of intervals and the number of clustering bins were positively correlated with the final number of nodes with (*r* = 0.32) and (*r* = 0.6), respectively whereas the overlap percentage was found to be negatively correlated with the final number of nodes (*r* = − 0.16). The average node size (average content in samples of each node) presented a negative correlation with the initial number of intervals (*r* = −0.4) and the number of bins used for clustering (*r* = −0.35) and positive correlations with the overlap percentage (*r* = 0.57). Finally, the number of ramifications in the graph was positively correlated with the initial number of intervals (*r* = 0.24) and the overlap percentage (*r* = 0.29) and was found to be negatively correlated with the number of bins (*r* = −0.35). Figure 1 A depicts the correlations between the aforementioned pairs of input and output parameters whereas Figure 1 B shows the combinatorial graph for a selection of parameter settings.

**Figure 1:**
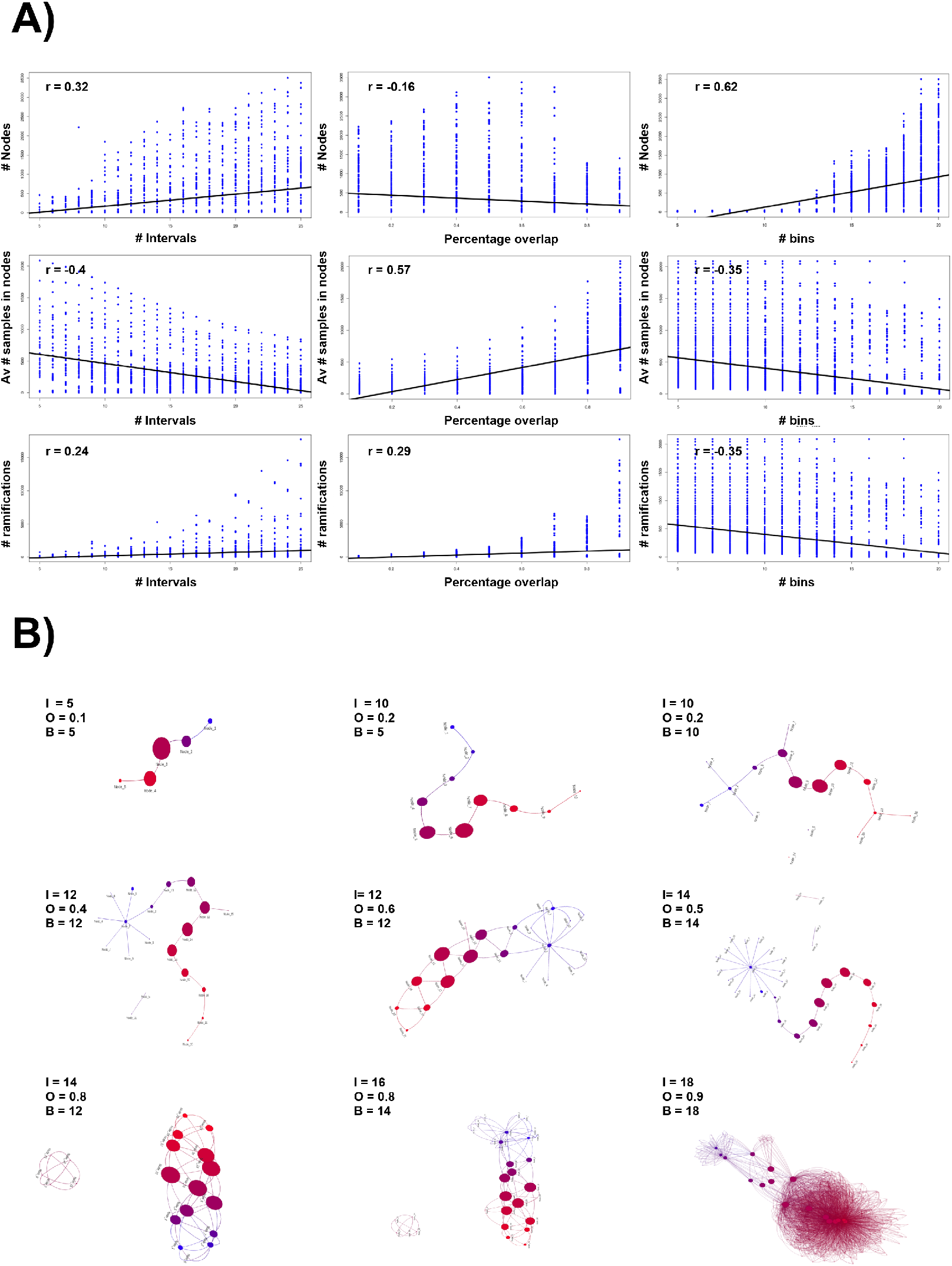
A) Correlations between input parameters (number of intervals, overlap percentages, and number of bins) and output features (number of nodes, average number of samples in nodes, and number of ramifications). B) Visualization of the SurvMap output results for breast cancer subset A under different parameter settings. Node hues represent the range of average values of the filtering function with blue hues indicating lower values and red hues depicting high values.

#### 3.2.2 Breast cancer subset A analyses results

Next, we ran SurvMap using the breast cancer subset A selecting the 200 genes identified by our feature selection criterion. Supplementary table 2 shows the list of genes retrieved by the **gene selection surv()** function with the **top bottom** option selected. The filter function was run using parameters *p* = 2 and *k* = 1. One dimensional mapper analysis was then carried out employing 8 initial intervals, an overlap percentage of 0.5, 8 bins, hierarchical clustering, and euclidean distances. SurvMap produced a combinatorial graph with 9 nodes with the following sample distribution (Node 0: 93, Node 1: 219, Node 2: 700, Node 3: 1261, Node 4: 1189, Node 5: 619, Node 6: 187, Node 7: 29, and Node 8: 1). Figure 2 A shows the combinatorial graph output generated by SurvMap. To carry out survival and differential expression analyses, promiscuous samples placed in more than one node were assigned to a unique node employing the **assign promiscuous samples()** function. After individual node assignment, 8 nodes presented more than 10 samples. (Node 0: 66, Node 1: 66, Node 2: 332, Node 3: 531, Node 4: 632, Node 5: 378, Node 6: 140, and Node 7: 27).

**Figure 2:**
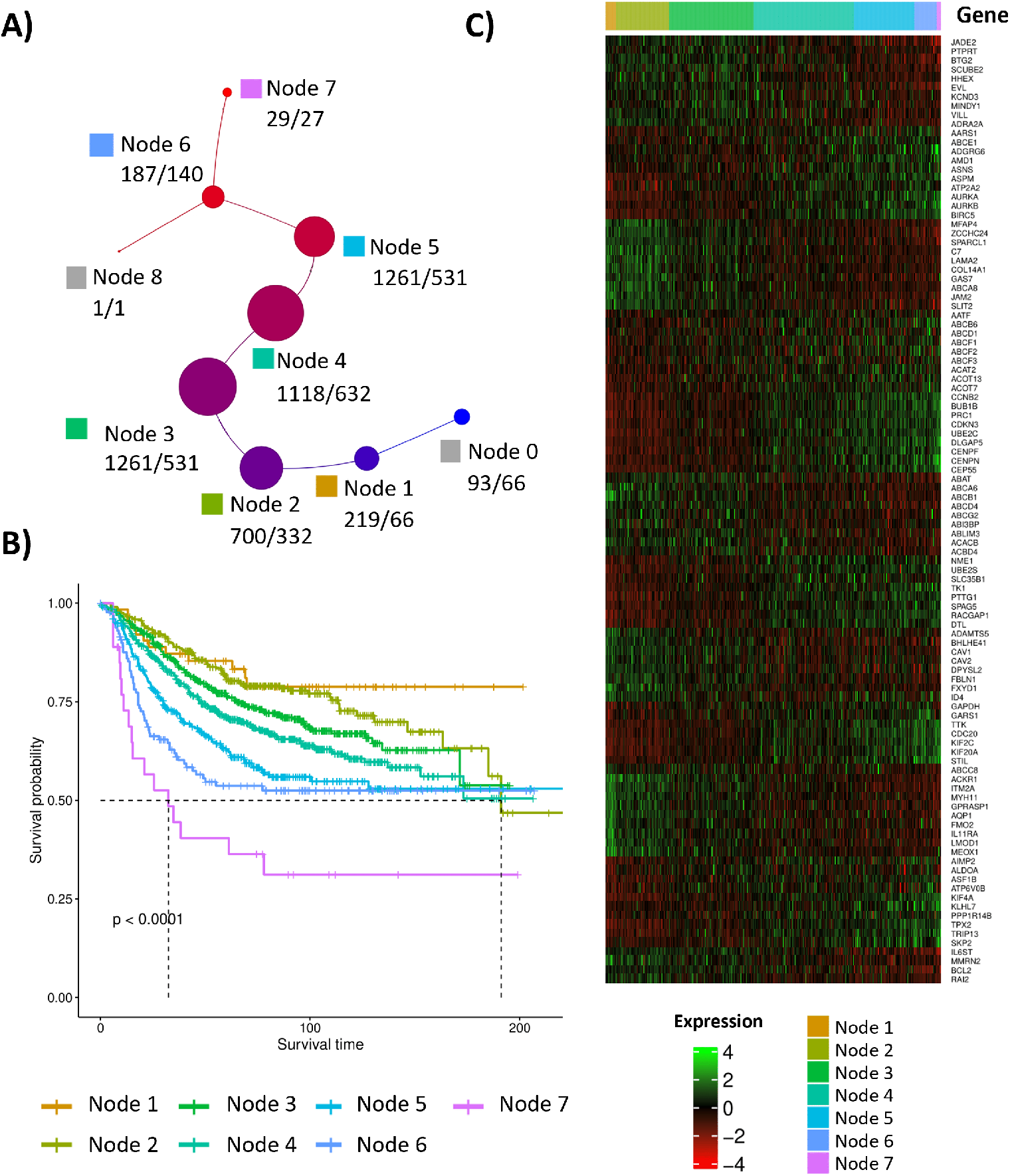
A) SurvMap output results for breast cancer subset A using parameter values of 10 for the number of intervals, 0.4 for the overlap percentage and 15 for the number of bins. B) Kapplan-Meier plots depicting disease free survival of those nodes that included more than 20 samples. C) Heatmap showing the top 5 up- and down-regulated genes identified in each node by performing Wilcoxon tests comparing the expression levels of the samples included in a particular node to all other samples.

Survival analysis (log-rank tests) showed significant differences in relapse-free survival among nodes (*p* = 8 × 10^−14^). Node 0 was not included in the survival analysis since all samples placed at this node were derived from healthy control tissues. Node 1 presented the best prognosis in terms of disease-free survival followed by node 2. Nodes 3, 4, and 5 presented shorter relapse-free survival times compared to Nodes 1 and 2 whereas nodes 6 and 7 contained the samples from those patients displaying the worse prognosis, indicating the ability of PAD-S and SurvMap to identify specific sets of samples with diverse clinical outcomes. Figure 2 B shows the Kaplan-Meier plot for identified nodes containing more than 10 samples.

Gene markers for each node were established through differential gene expression analyses (two tailed Wilcoxon tests) by comparing the samples assigned to a particular node to all other samples employing the **perform wilcoxon each()** function. Differential expression analyses were carried out employing the complete dataset A gene expression matrix. Supplementary File 4 includes the complete differential gene expression analysis results for each node. Table 2 shows ten of the most significantly up-regulated and down-regulated genes identified in each node and Figure 2 C displays a heatmap showing the expression levels of the node marker genes. Given the nature of SurvMap, changes in the expression levels of particular node markers present a smooth transition between consecutive nodes. The selection of the top differentially expressed genes for each node was based on their adjusted p-values. Genes with the same adjusted p-values where ordered alphabetically.

**Table 2:**
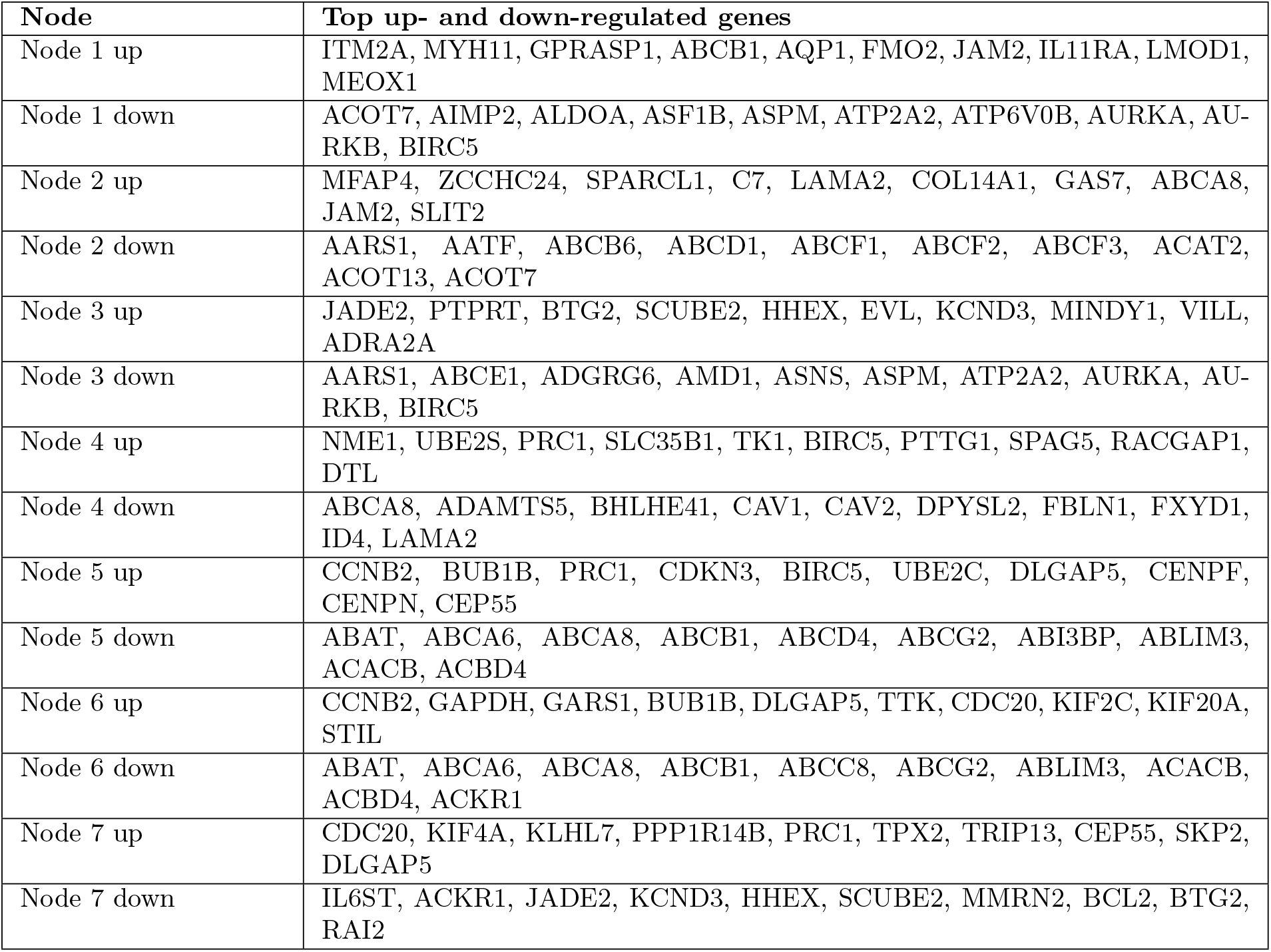
Top 10 up- and down-regulated genes in each node when compared to samples placed in all other nodes.

GSVA analysis results showed a transition in the gene expression levels of several hallmark pathways from good to poor prognosis nodes, including a gradual increase in the expression levels of G2M check-points, E2F and MYC targets, MTORC1 signaling, and genes involved in the glycolysis, and the response to unfolded proteins, among others. A gradual decrease in the expression levels of pathways linked to estrogen response, angiogenesis, and apoptosis was also observed. Figure 3A depicts the average GSVA scores for each Hallmark’s pathway observed in each node.

**Figure 3:**
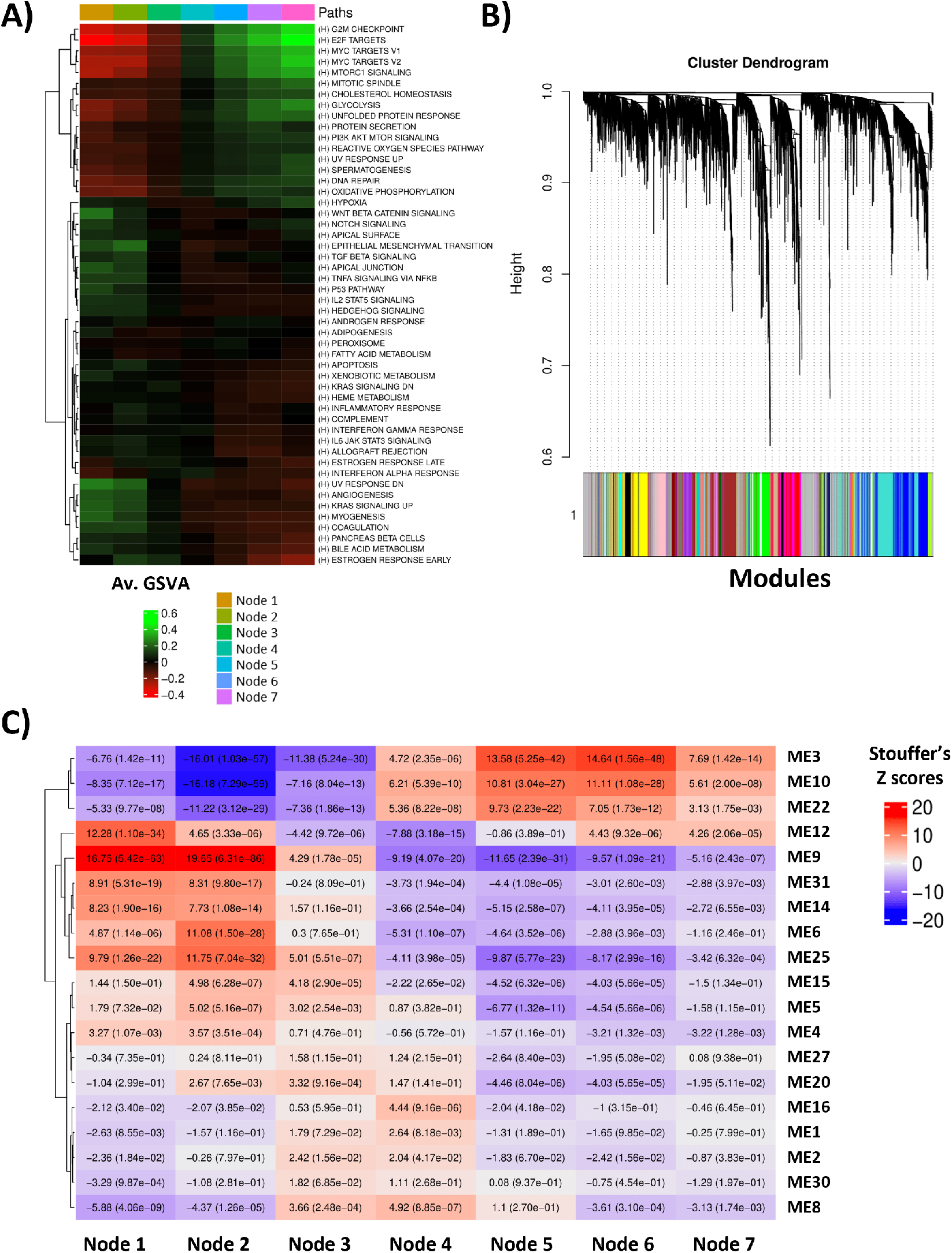
A) GSVA analysis results. The heatmap represents the average values of the GSVA score observed in each node for each tested pathway. Green values indicated over-expressed pathways whereas red values are employed for down-regulated pathways. B) Dendrogram obtained by WGCNA depicting the identified modules of co-expressed genes. C) Measures of association between nodes and co-expression modules eigengens obtained combining the results of each dataset through Stouffer’s method. Red hues depict positive measures of association whereas blue hues display negative associations.

Next, we assessed how several known covariates were distributed in each node. First, we analyzed the distribution of the PAM50 intrinsic subtypes identified by the genefu package. Node 1 was composed mostly of samples classified as Luminal A (46%) and normal-like (50.79%). The most represented intrinsic subgroup in node 2 was luminal A (79.11%) followed by samples classified as normal-like (10.12%), and luminal B (5.38%). In the case of node 3, 48.57% of samples were luminal A, a 25.82% luminal B, and a 14.75% basal.

Node 4 presented the highest proportion of luminal B samples (45.78%) followed by basal-like samples (22.37%). Her2 samples were mostly distributed across nodes 5, 6, and 7 representing 20.48%, 26.71%, and 25.93% of samples respectively. Finally, the highest proportion of basal-like samples was observed in nodes 6 and 7 which contained 43.51%, and a 51.85% of basal-like samples respectively. Therefore, despite nodes tend to be over-represented with specific intrinsic sub-types all they are composed of a mixture of them indicating that mapper highlights different aspects of the underlying biology of breast cancer.

Tumor grade was found to gradually increase when transitioning from good to poor prognosis groups with node 1 presenting 32.14% of samples derived from grade 1 tumours and 28.57% of grade 3 tumors and nodes 6 and 7 presenting around 76% of grade 3 tumours.

Node 1 samples were derived from patients with the lowest percentage of affected lymphatic nodes (36.96%) whereas node 7 presented the highest percentage (52.38%). And the lowest percentages of ER and PR positive samples were observed in nodes 5 and 6. Supplementary Tables 3-7 show the distribution of the collected co-variates in each node.

Weighted Gene Co-expression analysis was carried out as follows. First, the powers that generated adjacency matrices presenting the best scale-free topology fit were selected using the *pickSoftThreshold* function using parameters *bicor* for the correlation function and signed hybrid networks. The powers selected for each dataset can be found in Supplementary Table 8 whereas Supplementary Figures 1 and 2 show the scale-free topology fit and mean connectivities plotted against the soft threshold powers for each dataset, respectively. Next consensus module analysis was carried out employing the *blockwiseConsensus-Modules* function with parameters mergeCutHeight = 0.35, deepSplit = 2, and consensusQuantile = 0.25. Thirty-nine consensus co-expression modules, including the ”grey” or zero module, were identified by the analysis (See Figure 3 B).

Modules were tested for enrichment in specific biological processes from Gene Ontology (GO). Nineteen modules were found to be significantly enriched in different GO terms. Supplementary File 5 shows the enrichment analysis results for genes placed in each co-expression module. Then, we determined whether specific modules were associated with the nodes identified by SurvMap by computing correlations between node membership status and the module eigengenes computed by the *multiSetMEs* function. The individual results for each dataset were combined meta-analytically using Stouffer’s method. Several co-expression modules were found to be positively associated with the good prognosis nodes while presenting negative associations with the worse prognosis nodes. Those included consensus ME12 which was found to be enriched in biological processes linked to epidermal cell differentiation (p-adj = 9.4 × 10^−6^), canonical Wnt signalling (p-adj = 7.7 × 10^−6^), and keratinization (p-adj = 6.0 × 10^−5^), ME9 which was enriched in genes linked to angiogenesis (p-adj = 1.6 × 10^−17^), epithelial cell proliferation (p-adj = 2.0 × 10^−12^), cell adhesion (p-adj = 6.3 × 10^−8^), and fat cell differentiation (p-adj = 7.2 ×10^−7^), ME31 that was enriched in genes linked to ribosome biogenesis (p-adj = 3.2 × 10^−22^) and RNA processing (p-adj = 7.4 × 10^−11^), ME14 that was linked to RNA metabolic process (p-adj = 6.6 × 10^−12^) and RNA splicing (p-adj = 7.2 × 10^−12^), and ME6 which was found to be enriched in angiogenesis (p-adj = 2.0 × 10^−24^), cell motility (p-adj = 2.5 × 10^−22^), and cell-matrix adhesion (p-adj = 2.4 × 10^−13^) related genes. In contrast, several co-expression modules presented negative associations with the better prognosis nodes and positive associations with worse prognosis nodes. Instances of the latter include ME3, ME10, and ME22. ME3 was found to be enriched in genes linked to DNA repair (p-adj = 5.8 × 10^−27^), nuclear division (p-adj = 8.4 × 10^−27^), mitotic nuclear division (p-adj = 4.2 × 10^−25^), DNA replication initiation (p-adj = 5.4 × 10^−20^), and cell cycle checkpoint signalling (p-adj = 5.3 × 10^−17^) among others. ME10 was enriched in ATP metabolic process (p-adj = 1.7 × 10^−11^), oxidative phosphorylation related genes (p-adj = 8.5 × 10^−9^), as well as protein folding (p-adj = 5.5 × 10^−8^). ME22 was related to genes involved in translation (p-adj = 2.4 × 10^−8^). Other co-expression modules presented more complex patters of association with the nodes detected by PAD-S and SurvMap. For instance ME12 was found to be positively associated with Nodes 1 and 2, negatively associated with nodes 3 and 4 and positively associated with nodes 6 and 7. ME12 was found to be enriched in genes related to epidermis development (p-adj = 1.5 × 10^−8^), cytoskeleton organization (p-adj = 1.0 × 10^−6^), regulation of Wnt signalling pathway (p-adj = 2.4 × 10^−6^), and extracellular matrix organization (p-adj = 6.8 × 10^−5^). Figure 3 C shows the results of the analysis of associations between co-expression modules and nodes.

#### 3.2.3 Breast cancer subset B analysis and subsets A and B analyses results comparison

Then, we repeated the analyses using subset B employing the same configuration of parameters as the one utilized in the analysis of subset A. The same 200 genes as those employed in dataset A analysis were also selected. Subset B included 2187, divided into 2115 tumour samples and 72 healthy tissues. PAD-S and SurvMap identified 8 nodes (Nodes 0 to 7) including 105, 291, 830, 1214, 1082, 586, 167, and 36 samples respectively. After individual node assignment, the number of samples placed in each node was 70, 143, 333, 484, 579, 414, 132, and 32. (See Supplementary Figure 3). Significant differences in relapse-free survival among nodes were also identified (p-val = 5 × 10^−14^), with an increasing worse prognosis when moving from Node 1 to Node 7. Supplementary Figure 4 shows the Kaplan-Meier curves for nodes identified in subset B.

To determine whether there was a correspondence between the nodes identified employing subsets A and B, we compared the outputs of their differential gene expression analysis results by calculating overlap coefficients between all pairwise combinations of the nodes identified in each subset. We also computed correlations between the average gene expression profiles of the samples placed in each node. The differential gene expression analysis results for subset B can be found in Supplementary File 6. The overlap coefficient analyses showed similarities in terms of the patters of differential gene expression observed in both subsets. In general, nodes containing samples derived from patients with longer relapse-free survival times tended to present similar patterns of differential gene expression. For instance, the node 1 identified in dataset A presented overlap coefficients of 0.75 and 0.71 for up-regulated and down-regulated genes when compared to dataset B node 1 and samples included in nodes 2 of datasets A and B presented overlap coefficients of 0.85 and 0.81 for up-regulated and down-regulated genes, respectively. In summary, node 1 from dataset A presented the highest overlap coefficients for both up- and down-regulated genes with node 1 from dataset B. Node 2 from dataset A did it with node 2 of dataset B, the same pattern was observed for nodes 6, and 7. Node 5 from dataset A presented the highest overlap coefficients with node 6 from dataset B and similar overlap coefficients with node 5 from dataset B. Figure 4 A and B shows the overlap coefficients of the up- and down-regulated genes for all pair-wise node combinations between subsets A and B. Similar trends were observed in the correlations analysis comparing the average gene expression profiles of all pair-wise combinations of nodes from each dataset with nodes with similar prognoses presenting higher correlations. (See Figure 4 C). In general, these results suggest that SurvMap and PAD-S are able to identify groups of samples with compatible relapse-free survivals and similar transcriptomics profiles in two independent cohorts.

**Figure 4:**
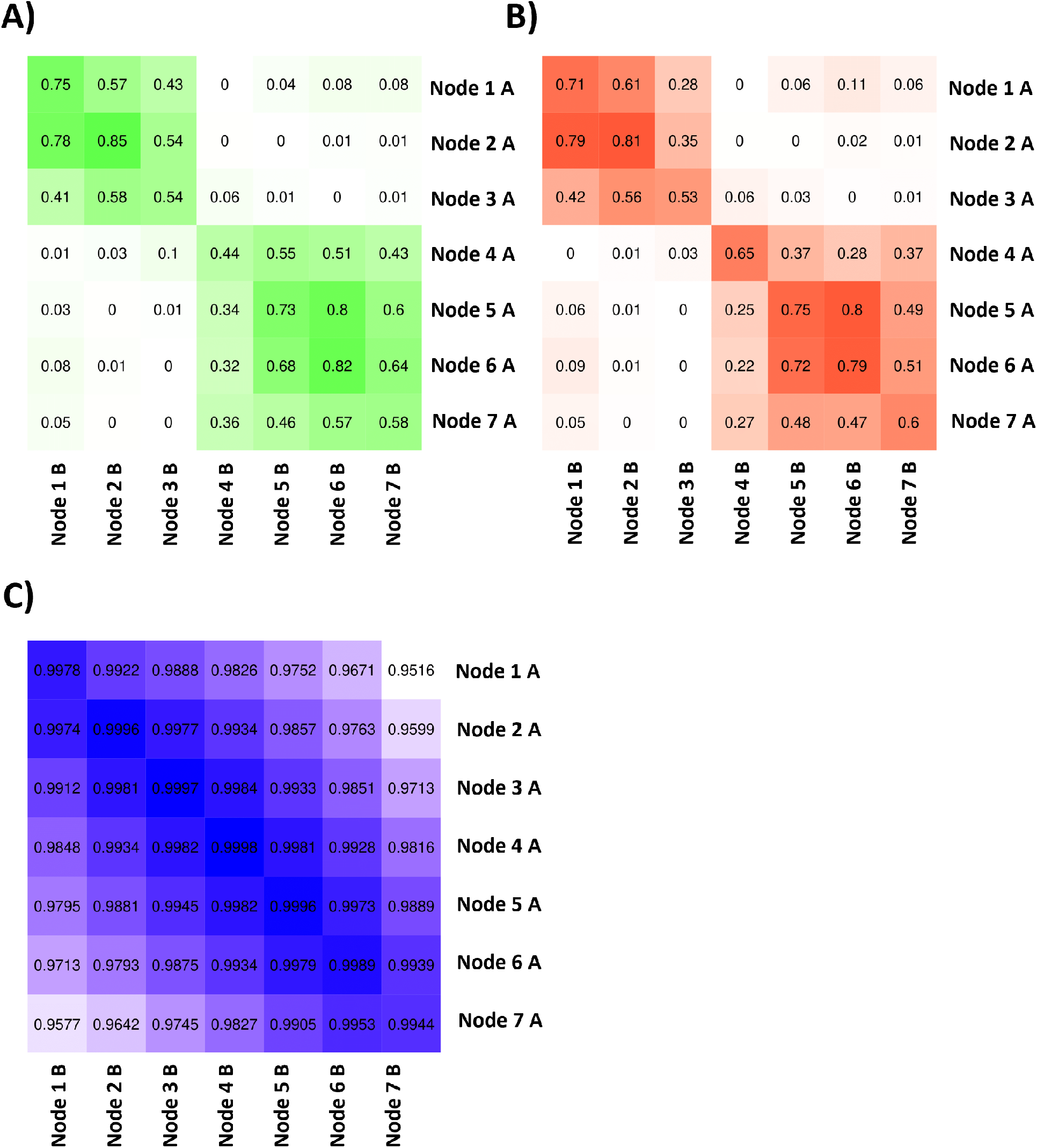
A) Overlap coefficients for the genes jointly up-regulated in all pairwise combinatios of datasets A and B nodes identified by PAD-S and SurvMap. B) Overlap coefficients for the genes jointly down-regulated in all pairwise combinatios of datasets A and B nodes identified by PAD-S and SurvMap. C) Correlations between the gene expression profiles of ll pairwise combinations of datasets A and B nodes identified by PAD-S and SurvMap.

#### 3.2.4 Using SurvMap to identify subgroups of samples inside canonical breast cancer groups (PAM50)

Next, we used the complete breast cancer dataset to test whether PAD-S and SurvMap were able to identify subgroups of samples with significantly different relapse times and molecular profiles inside the breast cancer intrinsic subtypes (Lumminal A, Lumminal B, Basal, and Her2) identified by the PAM50 algorithm implemented in the genefu package. The complete dataset included 1378 samples identified as luminal A. SurvMap was ran using 4 initial intervals, p = 0.5, euclidean distances, hierarchical clustering, and the standard mode for optimal number of clusters identification with 8 bins. Using this setup, we were able to identify 4 different nodes. After promiscuous sample assignment to individual nodes, they included 417, 541, 420, and 141 samples respectively. Significant differences in relapse-free survival were observed between these nodes using log-rank tests (p-val 1 × 10^−7^) (Figure 5 A). Several genes were found to be unregulated in better prognosis nodes compared with worse prognosis nodes including *PDGFRA, C1S, MFAP4, NDN*, and *ZCCHC24* among many others, whereas other genes were found to present lower expression levels in better prognosis compared to worse prognosis nodes including *P4HTM, SPR, NSF, ATP6AP1*, and *DHPS*. In terms of hallmark pathways the better prognosis groups and particularly node 1 was characterized by lower expression levels of pathways linked proliferation such as E2F targets and G2M checkpoints, as well as MTOR signalling, DNA repair and MYC targets. Worse prognosis nodes inside luminal A samples presented higher expression levels of the previous pathways. In contrast Luminal A node 1 was characterized by the presence of higher expression levels of angiogenesis, epithelial mesenchymal transition, apoptosis, and immune system related pathways compared with worse prognosis luminal A nodes. (See Supplementary Figure 5). Supplementary File 7 presents the differential gene expression analysis results for luminal A nodes.

**Figure 5:**
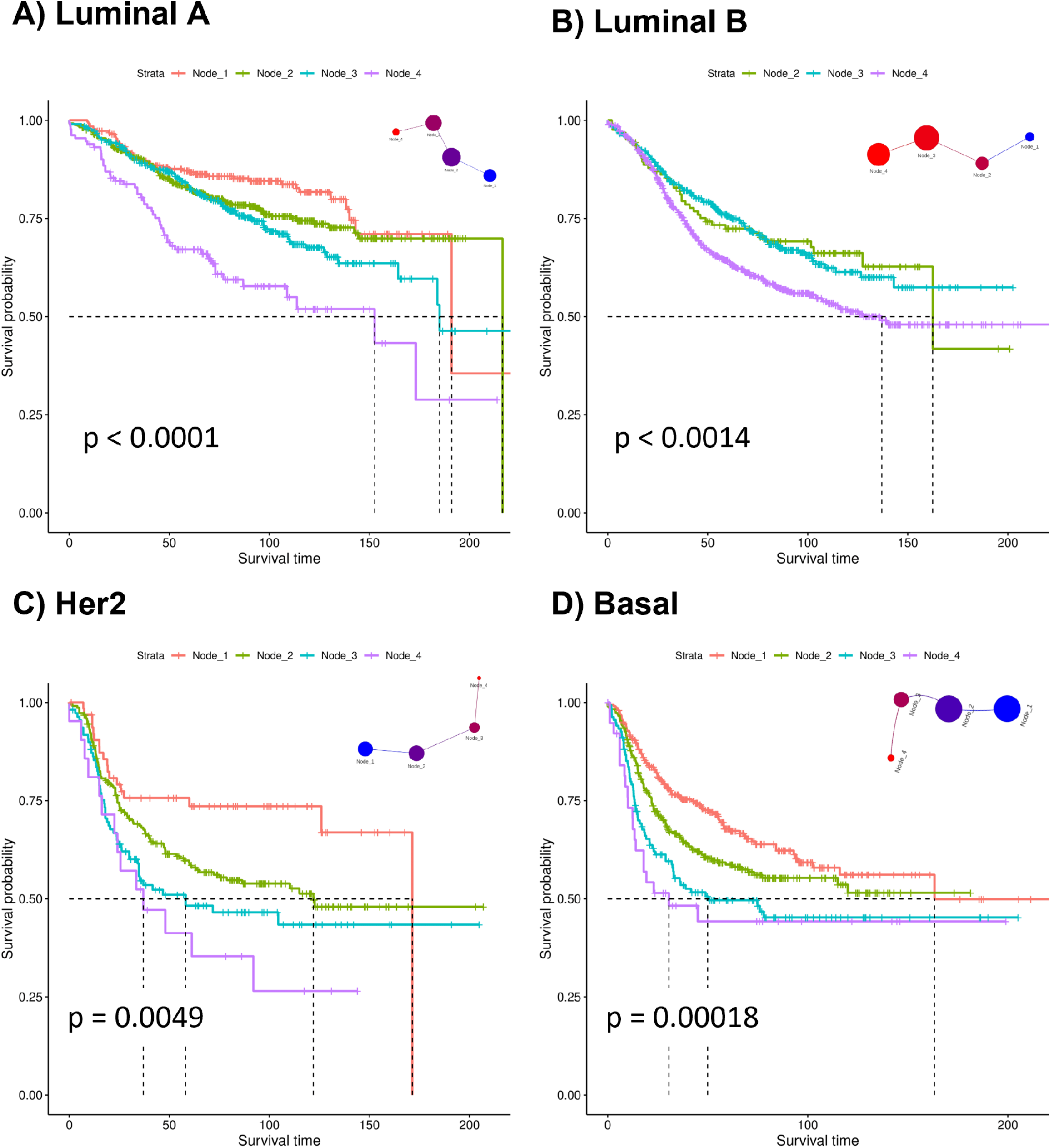
Kapplan-Meyer curves for the nodes identified by PAD-S and SurvMap insade each breast cancer intrinsic subtype. A) Luminal A, B) Luminal B. C) Her2. D) Basal.

One thousand two-hundred and eighty three samples were classified as luminal B samples by the genefu package. SurvMap produced 4 nodes that included 140, 138, 420, and 726 samples, respectively. Node 1 was entirely composed of samples derived from healthy breast tissues and was therefore not included in survival analysis. We observed significant differences in terms of disease-free survival between the three remaining nodes (p-val = 0.001) (Figure 5 B). Genes with higher expression levels in better prognosis groups included *PRKCB, CCR7, LCK, TRBC1*, and *TRAC*, among others, wheres genes presenting lower expression levels in better prognosis nodes compared to worse prognosis nodes in luminal B samples included *EEF1A2, ENO2, HSPB1, P3H4*, and *P4HA2* among others. Better prognosis nodes in luminal B samples presented higher expression levels of immune related processes, such as interferon alpha and gamma responses, allograft rejection, inflammatory response, and IL6 JAK STAT3. Better prognosis groups also present lower expression levels of genes linked to the oxidative phosphorylation, and the glycolysis, among others. (See Supplementary Figure 6). Supplementary File 8 presents the differential gene expression analysis results for luminal A nodes.

Our combined dataset contained 479 samples tagged as Her2 by the PAM50 algorithm. SurvMap identified 4 nodes including 217, 252, 127, and 24 samples respectively that presented significant differences in terms of disease-free survival (p-val = 0.005)(Figure 5 C). Genes up-regulated in better prognosis groups included *CCR7, LTB, CCL19, SELL*, and *CD247* among others. In contrast, *ACKR1, ADA2, AIF1, AOAH*, and *ARHGAP25* were found to be up-regulated in worse prognosis nodes compared with worse prognosis nodes. Her2 samples placed at worse prognosis nodes presented increased expression levels E2F targets, G2M checkpoints, MYC targets, as well as MTORC1 signaling, DNA repair and unfolded protein response among others and lower expression levels of IL6 JACK STAT3 signaling, allograft rejection, and fatty acid metabolism, among others. (See Supplementary Figure 7). Supplementary File 9 presents the differential gene expression analysis results for basal nodes.

The dataset contained 858 samples assigned to the basal group. In this particular case we selected parameters p = 4 and k = 2 as parameters for the filtering function. Four nodes were identified inside basal-like samples presenting significant differences in terms of relapse-free survival. (p-val = 2 × 10^−4^)(See Figure 5 D). Better prognosis nodes inside basal samples presented higher expression levels of genes such as *TRAC, CD2, CD3D, TRBC1*, and *CD27* compared to worse prognosis nodes whereas *PTDSS1, SLC19A1, BIRC5, DSCC1*, and *NT5DC2* presented lower expression levels in good prognosis nodes compared to the worse prognosis ones. The worse prognosis nodes presented higher expression levels of pathways linked to E2F targets, MYC target, and G2M checkpoints among others and lower expression levels linked to immune related processes, such as interferon alpha and gamma signaling, and apoptosis, among others. (See Supplementary Figure 8). Supplementary File 10 presents the differential gene expression analysis results for basal nodes.

Finally, 221 samples were classified as normal-like. SurvMap identified 4 nodes with no significant differences in terms of relapse-free survival (p-val = 0.1).

Supplementary Table 1 shows the intrinsic subtype classification of all samples included in the dataset.

### 3.3 Testing PAD-S and SurvMap using Breast Cancer Methylation Data

Finally, to demonstrate the ability of PAD-S and SurvMap to works with different data types other than transcriptomic data, we tested its performance in a breast cancer methylation dataset derived from TCGA. It included the methylation levels of 485577 probes derived from 895 samples of which 97 where healthy breast samples, 793 were primary breast tumors and 5 were metastatic samples. Probes presenting missing values in more than 50% of the samples were filtered out, leaving a total of 395777 probes. The healthy state model was then constructed using the 97 available healthy breast tissue samples and disease components were computed for the whole dataset. Cox proportional hazard models were fitted to measure the association of the methylation levels of each probe with overall survival. We selected the top 200 probes identified by our feature selection criteria. In this case we used the following setup: We selected an initial number of intervals of 4, an overlap percentage of 0.5, Pearson’s correlations as distance types, and silhouette as the optimal number of clusters assessment method. The values for parameters *p* and *k* for the filtering function were 2 and 1 respectively. PAD-S and SurvMap identified 6 nodes that after promiscuous sample assignment to individual nodes included 444, 281, 135, 2, 32, and 1 samples for nodes 1, 2, 3, 4, 5, and 6, respectively (Figure 6 A). Highly significant differences in overall survival were observed between nodes (P ≤ 2*e* − 16) (Figure 6 B). With Node 1 being the best prognosis node in terms of overall survival and node 5 the worst. Differential methylation analyses between the samples included in a particular node compared to those that were placed in all other nodes were carried out. Table 3 shows the top 5 probes associated to genes that were found to be differentially methylated between groups. Supplementary File 11 shows the differential methylation analysis results. Enrichment analysis was carried out using the *gometh* function from the missMethyl package. Probes displaying high methylation levels in node 1 were enriched in phosphorus metabolic processes (p-ajd = 9.64 × 10^−9^), cytoskeletal protein binding (p-adj = 3.62 × 10^−5^), and protein kinase activity (p-adj = 6.19 × 10^−04^), among others, whereas probes with lower methylation levels in node 1 compared to all other groups were found to be enriched in processes linked to cell adhesion (p-adj = 4.89 × 10^−14^) and DNA-binding transcription factor activity (p-adj = 3.51 × 10^−11^). Worse prognosis nodes (node 3) presented higher methylation levels of probes associated to genes linked to cell-cell signaling, (p-adj = 9.47 × 10^−24^), cell adhesion (p-adj = 8.21 × 10^−22^), leukocyte activation (p-adj = 1.01× 10^−5^), and angiogenesis (p-adj = 2.66 × 10^−5^) among others and lower levels of methylation in probes linked to genes associated with cytoskeletal protein binding (p-adj = 2.95 × 10^−6^). Supplementary File 12 shows the enrichment analysis results for differentially methylated probes.

**Figure 6:**
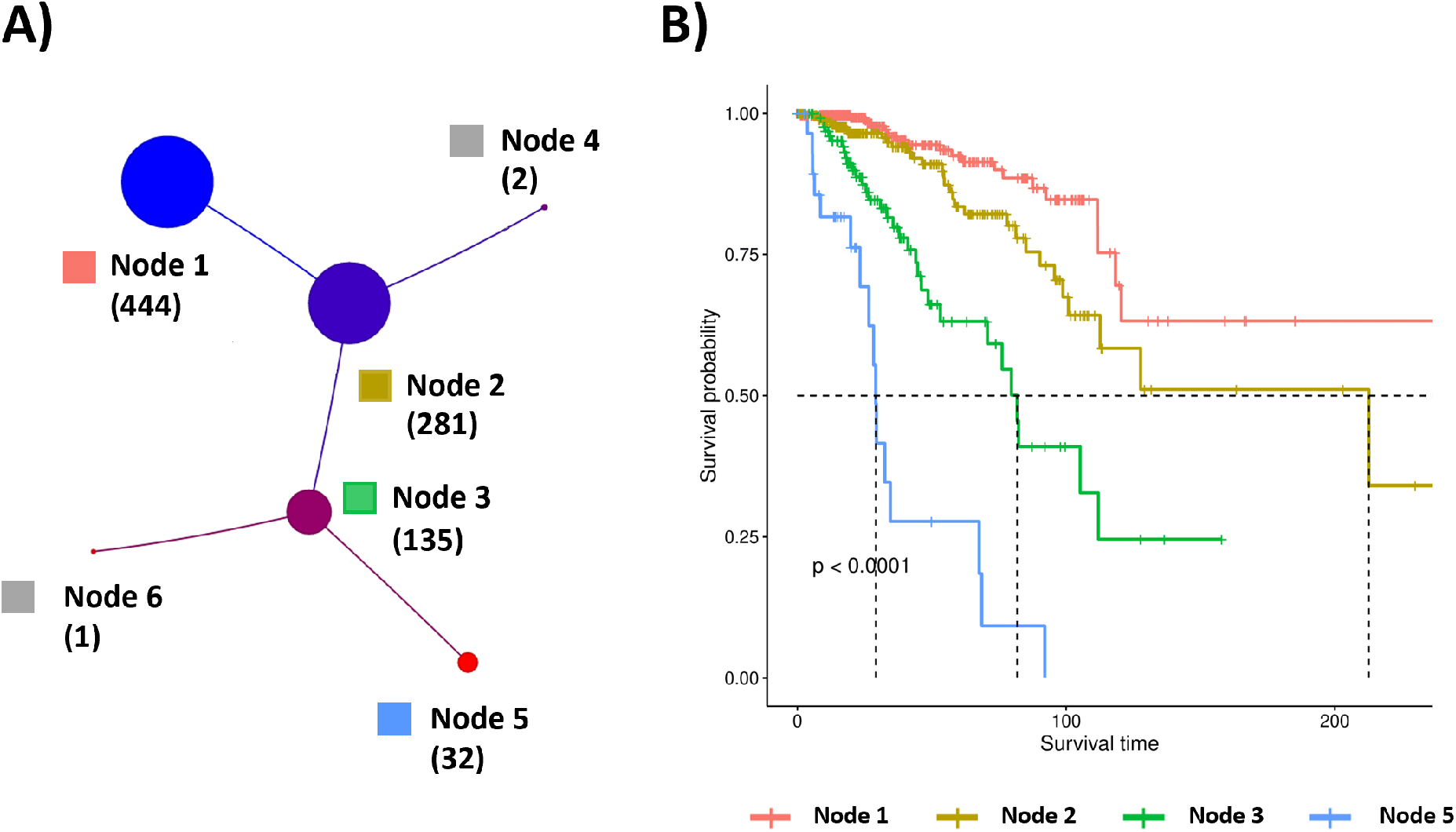
A) Nodes detected by PAD-S and SurvMap in breast cancer methylation data derived from TCGA. B) Kaplan-Meier depicting the overall survival probabilities of patients included in nodes 1, 2, 3, and 5.

**Table 3:**
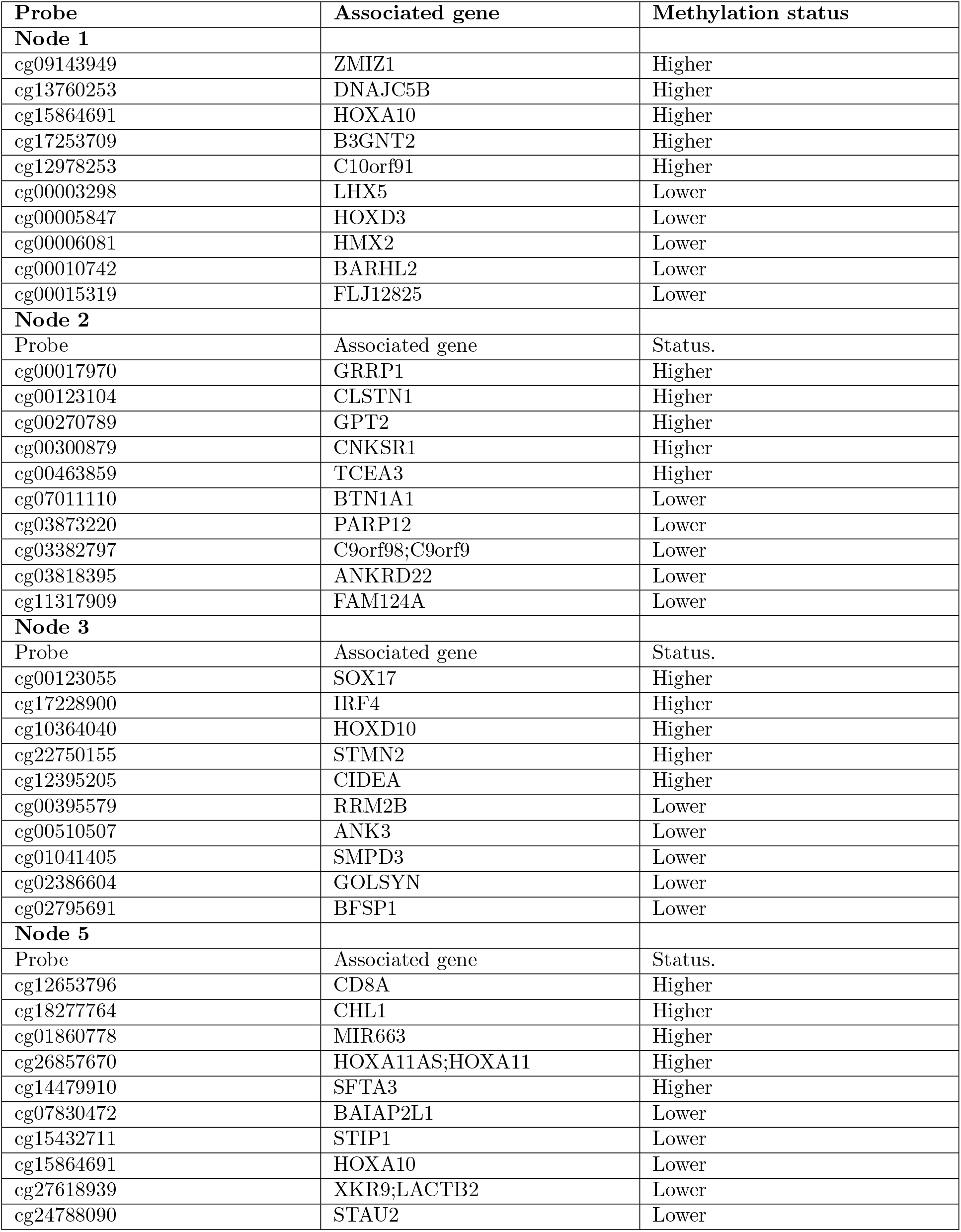
Top 5 hyper- and hypo-methylated probes in the samples placed in a particular node when compared to samples placed in all other nodes.

## 4 Discussion

Here we presented Progression Analysis of Disease with Survival (PAD-S). A method that extends PAD analysis by integrating information about the degree of association of each feature with survival through the use of specific feature selection procedures and a new adapted filter function. In addition, we also integrated a novel method for Gaussian noise removal applied to the construction of the healthy state model, as well as other improvements such as an option to select a Silhouette-based method to recover the optimal number of clusters when running the Mapper algorithm, as well as a strategy to place promiscuous samples into unique nodes based on a singular value decomposition approach, that allows downstream analysis such as differential gene expression and survival analysis. We implemented PAD-S in an R package called SurvMap that provides the user with the necessary functions to run both PAD-S and traditional PAD analysis. Auxiliary functions aimed to carry out differential expression and single sample enrichment analysis were also integrated into the package.

PAD-S and SruvMap were able to identify nodes containing samples with idiosyncratic gene expression and pathway signatures and distinct behaviors in terms of disease-free survival. Several genes found to be up-regulated in the best prognosis node (Node 2) have been previously linked to a good prognosis in breast cancer. For instance, high expression levels of *JAM2*, which encodes for a junctional adhesion molecule of the immunoglobulin family, are associated with a better prognosis in breast cancer [20]. Similar patterns of association with prognosis have been reported for the Integral Membrane Protein 2A tumor suppressor gene *ITM2A* [21] and for the gene encoding for the myosin heavy chain 11 (*MYH11*). which has been linked to better clinical outcomes in terms of both overall and disease-free survival in breast cancer [22]. In addition, the loss of constitutive *ABCB1* expression, one of the top up-regulated genes in node 2, has been linked to worse prognosis in breast cancer patients [23] and breast cancer patients with high expression levels of *AQP1* treated with anthracyclines are known to display better clinical outcomes relative to those with low *AQP1* levels [24]. Opposite patterns of association with prognosis have been reported for genes found to be down-regulated in node 2, including *AURKA* for which high levels of expression are linked to worse prognosis in estrogen receptor-positive breast cancers [25], *BIRC5* gene, which encodes for survivin know as the cell death preventing protein for which expression levels correlate with poor survival in different cancer types, including breast cancer [26].

Increased expression levels of several genes up-regulated in the nodes displaying the worst prognosis (Nodes 7 and 8) have also been previously linked to disease progression in breast cancer. *CCNB2* has been associated with lymphovascular invasion and high expression levels of this gene are linked to a worse prognosis [27, 28]. The same kind of associations with survival have been reported for the disc large-associated protein gene *DLGAP5* [29], the spindle assembly checkpoint kinase tyrosine/threonine kinase *TTK* [30], *CDC20* [31], *KIF20A* [32], *PRC1* [33], and *TPX2* [34]. For all those genes high expression levels have been correlated with poor clinical behavior in breast cancer.

In addition, genes down-regulated among poor prognosis nodes included *ACKR1, HHEX, SCUBE2, BCL2, BTG2*, and *RAI2*, among others. Tumors with high expression levels of the atypical chemokine receptor 1 (*ACKR1*) present significantly longer relapse-free and overall survival than individuals with low expression levels of this gene [35]. Low levels of the transcription factor hematopoietically expressed homeobox (*HHEX*) are also related to poor prognosis in breast cancer [36]. Surprisingly, low expression levels of *BCL2* which plays a pivotal role in the regulation of cellular apoptosis and which over-expression inhibits apoptotic cell death and activates cellular proliferation and tumor progression, have been previously linked to worse outcomes in terms of disease-free survival. Therefore, a dual role of *BCL2* as both an oncogenic and tumor-suppressor gene has been proposed [37]. The up-regulation of the cell cycle arrest protein *BTG2* correlates with increased overall survival in breast cancer [38] and low expression levels of Retinoic acid-induced 2 (*RAI2*) have also been reported to correlate with shorter disease-free survival and overall survival in breast cancer [39].

Therefore, many of the top differentially expressed genes in nodes produced by PAD-S and SurvMap have been previously involved in breast cancer prognosis in a way consistent with their placement in nodes characterized by specific behaviors in terms of survival.

The analysis outputs from subset B substantially agreed with those derived from subset A in terms of survival, the differentially expressed genes found in each node, and the average correlations of their expression profiles. This highlights the capacity of PAD-S and SurvMap to recognize consistent sets of samples across datasets, which are characterized by similar behaviors in terms of prognosis and gene expression.

Furthermore, we were able to identify sets of samples with significantly different relapse-free survival times inside breast cancer intrinsic subgroups that presented characteristic gene expression profiles and diverse patterns of pathway activation. For instance, the best prognosis group identified inside luminal A samples presented low expression levels of genes linked to diverse biological processes including cell cycle checkpoints, MTORC1 signaling unfolded protein response, and DNA repair compared to the works luminal A prognosis node. In contrast, lower expression levels of apoptotic and immune system-related pathways were observed in the worse prognosis luminal A nodes compared to those presenting better clinical outcomes. PAD-S and SurvMap were able to pinpoint sets of nodes with diverse clinical outcomes in all major intrinsic subtypes (luminal A, luminal B, Her2, and Basal) except for samples tagged as normal-like demonstrating their ability to further refine previous sample classification schemes by exploring different aspects of the underlying biological features of samples.

PAD-S and SurvMap are a general-purpose methods that can be applied to a wide range of omics data, including mRNA, and miRNA expression derive from both array and RNAseq-based studies, methylation, and copy number, as long as the datasets of interest contain continuous variables. To demonstrate the capacity of PAD-S to operate using different omic data types we ran SurvMap on a large breast cancer methylation dataset derived from TCGA for which overall survival information was available. We identified four nodes of breast cancer samples with significantly different outcomes in terms of overall survival and methylation profiles.

Several methylation probes presenting higher methylation values in good prognosis groups compared to the poor prognosis ones have previously been associated with cancer survival. For instance, the intragenic DNA methylation of *ZMIZ1* has been found to serve as a survival predictor for different types of cancer including glioblastomas, astrocytomas, bladder cancer, and renal carcinomas, with higher methylation being associated with better prognosis [40], the hypomethylation of *B3GNT5* promoter (one of the genes liked to probes presenting high methylation levels in good prognosis nodes) has been described to contribute to B3GNT5 overexpression and to promote breast cancer aggressiveness [41].

Probes displaying low methylation levels in good prognosis nodes included cg00003298 and cg00010742 among many others. cg00003298 is associated with the *LHX5* gene. Methylation of islands proximal to genes linked to cell fate commitment such as LHX5 has been reported to have prognostic values in breast cancer [42], cg00010742 is linked to gene *BARHL2*. High methylation levels of *BARHL2* have been found to correlate with tumor size, grade, and stage and poor prognosis in non-muscle-invasive bladder cancer [43] and this gene has been identified as a marker of late breast cancer carcinoma stage in murine models [44].

Several probes exhibiting high methylation levels in poor prognosis nodes (nodes 3 and 4) have been previously implicated in breast cancer progression. Some instances include cg00123055 linked to *SOX17*, which is highly methylated in primary breast tumors [45]. In addition, *SOX17* has been found to decrease proliferation and tumor formation in a Wnt/*β*-catenin-dependent fashion acting as a tumor suppressor [46] and cg18277764 which is associated with the *CHL1* gene, whose hypermethylation has also been identified as a biomarker of poor prognosis in breast cancer [47] or miR-663 (cg01860778), which expression levels caused by promoter hypomethylation have been linked to chemotherapy resistance in breast cancer [48].

Furthermore, PAD-S and SurvMap could be used to identify groups of samples with diverse behaviors in terms of relapse-free or overall survival by employing combined data from multiple omic platforms and a single underlying algorithm. The construction of such multi-omic classifiers based on PAD-S and SurvMap is out of the scope of this work and will be the focus of further research.

## Acknowledgments

This work was supported by ESI International Chair at CEU-UCH and by the Universidad CEU-UCH Research Programme.

## Credit contribution statement

**Jaume Forés-Martos:** Conceptualization, Methodology, Software, Writing, Original draft preparation **Beatriz Suay-García:** Reviewing and Editing, **Raquel Bosch-Romeu:** Reviewing and Editing **María Carmen Sanfeliu-Alonso:** Reviewing and Editing **Antonio Falcó:** Conceptualization, Methodology, Software, Supervision, Reviewing and Editing **Joan Climent:** Conceptualization, Methodology, Supervision, Reviewing and Editing

